# Reverse engineering neural networks to characterise their cost functions

**DOI:** 10.1101/654467

**Authors:** Takuya Isomura, Karl Friston

## Abstract

This work considers a class of biologically plausible cost functions for neural networks, where the same cost function is minimised by both neural activity and plasticity. We show that such cost functions can be cast as a variational bound on model evidence under an implicit generative model. Using generative models based on Markov decision processes (MDP), we show, analytically, that neural activity and plasticity perform Bayesian inference and learning, respectively, by maximising model evidence. Using mathematical and numerical analyses, we then confirm that biologically plausible cost functions—used in neural networks—correspond to variational free energy under some prior beliefs about the prevalence of latent states that generate inputs. These prior beliefs are determined by particular constants (i.e., thresholds) that define the cost function. This means that the Bayes optimal encoding of latent or hidden states is achieved when, and only when, the network’s implicit priors match the process that generates the inputs. Our results suggest that when a neural network minimises its cost function, it is implicitly minimising variational free energy under optimal or sub-optimal prior beliefs. This insight is potentially important because it suggests that any free parameter of a neural network’s cost function can itself be optimised—by minimisation with respect to variational free energy.

## 1. Introduction

Cost functions are ubiquitous in scientific fields that entail optimisation—including physics, chemistry, biology, engineering, and machine learning. Furthermore, any optimisation problem that can be specified using a cost function can be formulated as a gradient descent. In the neurosciences, this enables one to treat neuronal dynamics and plasticity as an optimisation process (Marr, 1969; Albus, 1971; Schultz et al., 1997; Sutton & Barto, 1998; Linsker, 1988; Brown et al., 2001). These examples highlight the importance of specifying a problem in terms of cost functions, from which neural and synaptic dynamics can be derived. In other words, cost functions provide a formal (i.e., normative) expression of the purpose of a neural network and prescribe the dynamics of that neural network. Crucially, once the cost function has been established and an initial condition has been selected, it is no longer necessary to solve the dynamics. Instead, one can characterise the neural network’s behaviour in terms of fixed points, basin of attraction, and structural stability—based on and only on the cost function. In short, it is important to identify the cost function to understand the dynamics, plasticity, and function of a neural network.

A ubiquitous cost function in neurobiology, theoretical biology, and machine learning is model evidence, or equivalently, marginal likelihood or surprise; namely, the probability of some inputs or data under a model of how those inputs were generated by unknown or hidden causes. Generally, the evaluation of surprise is intractable. However, this evaluation can be converted into an optimisation problem by inducing a variational bound on surprise. In machine learning, this is known as an evidence lower bound (ELBO), while the same quantity is known as variational free energy in statistical physics and theoretical neurobiology.

Variational free energy minimisation is a candidate principle that governs neuronal activity and synaptic plasticity (Friston et al., 2006; Friston, 2010). Here, surprise reflects the improbability of sensory inputs given a model of how those inputs were caused. In turn, minimising variational free energy, as a proxy for surprise, corresponds to inferring the (unobservable) causes of (observable) consequences. To the extent that biological systems minimise variational free energy, it is possible to say that they infer and learn the hidden states and parameters that generate their sensory inputs (Helmholtz, 1925; Knill & Pouget, 2004; DiCarlo et al., 2012) and consequently predict those inputs (Rao & Ballard, 1999; Friston, 2005). This is generally referred to as perceptual inference based upon an internal generative model about the external world (Dayan et al., 1995; George & Hawkins, 2009; Bastos et al., 2012).

Variational free energy minimisation provides a unified mathematical formulation of these inference and learning processes in terms of self-organising neural networks that function as Bayes optimal encoders. Moreover, organisms can use the same cost function to control their surrounding environment by sampling predicted (i.e., preferred) inputs. This is known as active inference (Friston et al., 2011). The ensuing free-energy principle suggests that active inference and learning are mediated by changes in neural activity, synaptic strengths, and the behaviour of an organism to minimise variational free energy, as a proxy for surprise. Crucially, variational free energy and model evidence rest upon a generative model of continuous or discrete hidden states. A number of recent studies have used Markov decision process (MDP) generative models to elaborate schemes that minimise variational free energy (Friston, FitzGerald et al., 2016; Friston, FitzGerald et al., 2017; Friston, Parr et al., 2017). This minimisation reproduces various interesting dynamics and behaviours of real neuronal networks and biological organisms. However, it remains to be established whether variational free energy minimisation is an apt explanation for any given neural network, as opposed to the optimisation of alternative cost functions.

In principle, any neural network that produces an output or a decision can be cast as performing some form of inference, in terms of Bayesian decision theory. On this reading, the complete class theorem suggests that any neural network can be regarded as performing Bayesian inference under some prior beliefs; therefore, it can be regarded as minimising variational free energy. The complete class theorem (Wald, 1947; Brown, 1981) states that for any pair of decisions and cost functions, there are some prior beliefs (implicit in the generative model) that render the decisions Bayes optimal. This suggests that it should be theoretically possible to identify an implicit generative model within any neural network architecture, which renders its cost function a variational free energy or ELBO. In what follows, we show that such identification is possible for a fairly canonical form of a neural network and a generic form of a generative model.

In brief, we adopt a reverse engineering approach to identify a plausible cost function for neural networks—and show that the resulting cost function is formally equivalent to variational free energy. Here, we define a cost function as a function of sensory input, neural activity, and synaptic strengths and suppose that neural activity and synaptic plasticity follows a gradient descent on the cost function. For simplicity, we consider single-layer feed-forward neural networks comprising firing rate neuron models and focus on blind source separation (BSS); namely, the problem of separating sensory inputs into multiple hidden sources or causes (Belouchrani et al., 1997; Cichocki et al., 2009; Comon & Jutten, 2010), which provides the minimum setup for modelling causal inference. Previously, we observed BSS performed by *in vitro* neural networks (Isomura et al., 2015) and reproduced this self-supervised process using an MDP and variational free energy minimisation (Isomura & Friston, 2018). These works suggest that variational free energy minimisation offers a plausible account of the empirical behaviour of *in vitro* networks.

In this work, we ask whether variational free energy minimisation can account for the normative behaviour of a canonical neural network that minimises its cost function, by considering all possible cost functions, within a generic class. Using mathematical analysis, we identify a class of cost functions—from which update rules for both neural activity and synaptic plasticity can be derived—when a single-layer feed-forward neural network comprises firing rate neurons whose firing intensity is determined by the sigmoid activation function. The gradient descent on the ensuing cost function naturally leads to Hebbian plasticity with an activity-dependent homeostatic term. We show that these cost functions are formally homologous to variational free energy under an MDP. Finally, we discuss the implications of this result for explaining the empirical behaviour of neuronal networks, in terms of free energy minimisation under particular prior beliefs.

## 2. Methods

In this section, we first derive the form of a variational free energy cost function under a specific generative model; namely a Markov decision process^1^. We will go through the derivations carefully, with a focus on the form of the ensuing Bayesian belief updating. The form of this update will re-emerge later, when reverse engineering the cost functions implicit in neural networks. This section starts with a description of Markov decision processes—as a general kind of generative model—and then considers the minimisation of variational free energy under these models.

### 2.1 Generative models

Under an MDP model (Fig. 1A), a minimal BSS setup (in a discrete-space) reduces to the likelihood mapping from *N*_*S*_ hidden sources or states 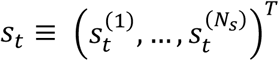 to *N*_*o*_ observations 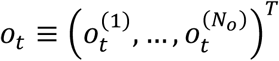. Each source and observation takes a value of one (ON state) or zero (OFF state) at each time step; i.e., 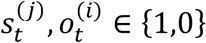. Throughout this paper, *j* denotes the *j*-th hidden state, while *i* denotes the *i*-th observation. The probability of 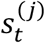 follows a categorical distribution 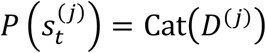, where 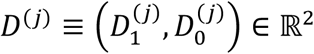 with 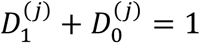.

**Figure 1.**
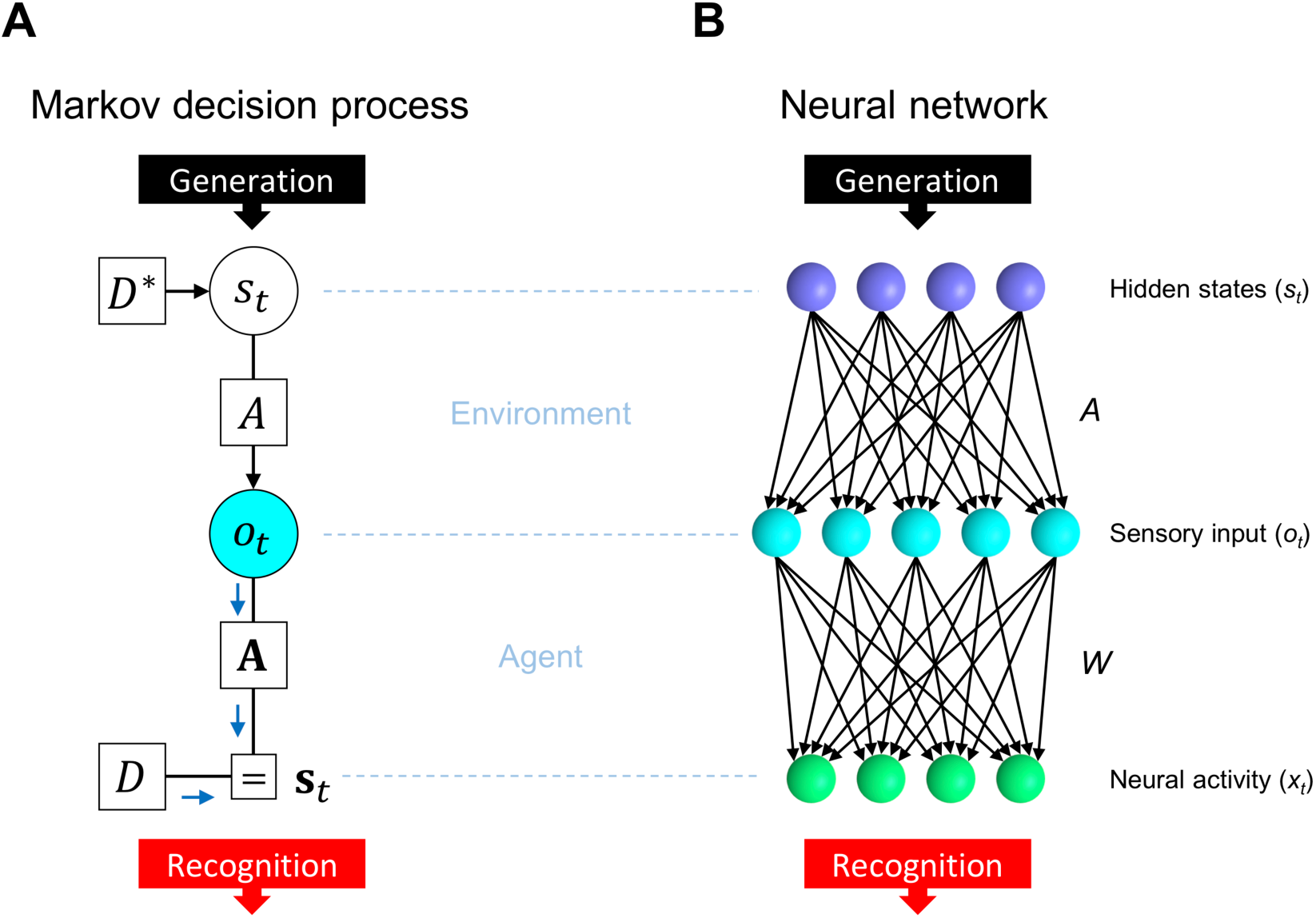
Comparison between an MDP scheme and a neural network. (**A**) MDP scheme expressed as a Forney factor graph (Forney, 2001; Dauwels, 2007) based upon the formulation in (Friston, Parr et al., 2017). In this BSS setup, the prior *D* determines hidden states *s*_*t*_, while *s*_*t*_ determines observation *o*_*t*_ through the likelihood mapping *A*. Inference corresponds to the inversion of this generative process. Here, *D*\* indicates the true prior while *D* indicates the prior under which the network operates. If *D* = *D*\*, the inference is optimal; otherwise, it is biased. (**B**) Neural network comprising a single layer feed-forward network with a sigmoid activation function. The network receives sensory inputs 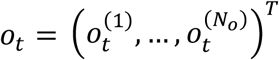 that are generated from hidden states 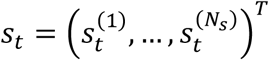 and outputs neural activities 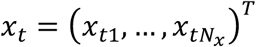. Here, *x*_*tj*_ should encode the posterior expectation about a binary state 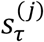.

The probability of an outcome is determined by the likelihood mapping from all hidden states to each kind of observation in terms of a categorical distribution, 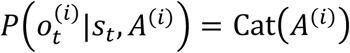. Here, each element of the tensor 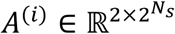 parameterises the probability that 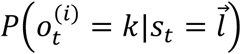, where *k* ∈ {1,0} are possible observations and 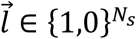 encodes a particular combination of hidden states. The prior distribution of each column of *A*^(*i*)^, denoted by 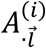, has a Dirichlet distribution 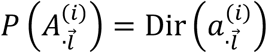 with concentration parameter 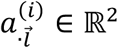. We use Dirichlet distributions, as they are tractable and widely used for random variables that take a continuous value between 0 and 1. Furthermore, learning the likelihood mapping leads to biologically plausible update rules, which have the form of associative or Hebbian plasticity: please see below and (Friston et al., 2016) for details.

We use *õ* ≡ (*o*_1_, …, *o*_*t*_) and 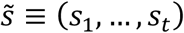 to denote sequences of observations and hidden states, respectively. With this notation in place, the generative model (i.e., the joint distribution over outcomes, hidden states, and the parameters of their likelihood mapping) can be expressed as

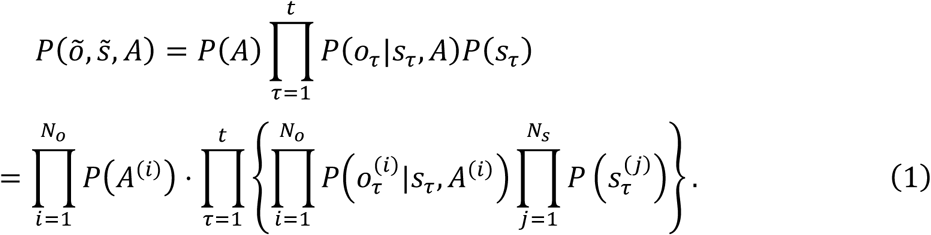

Throughout this paper, *t* denotes the current time, while *τ* denotes an arbitrary time from the past to the present, 1 ≤ *τ* ≤ *t*.

### 2.2 Minimisation of variational free energy

In this MDP scheme, the aim is to minimise surprise by minimising variational free energy as a proxy; i.e., performing approximate or variational Bayesian inference. From the generative model, we can motivate a mean-field approximation to the posterior (i.e., recognition) density as follows:

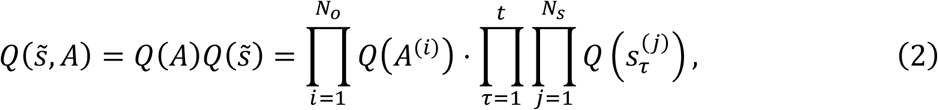

where *A*^(*i*)^ is the likelihood mapping (i.e., tensor), and the marginal posterior distributions of 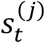 and *A*^(*i*)^ have a categorical 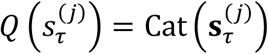 and Dirichlet distribution *Q*(*A*^(*i*)^) = Dir(**a**^(*i*)^), respectively. For simplicity, we assume that *A*^(*i*)^ factorises into the product of the likelihood mappings from the *j*-th hidden state to the *i*-th observation: 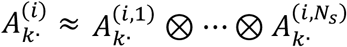 (where ⊗ denotes the outer product and *A*^(*i,j*)^ ∈ ℝ^2×2^). This (mean field) approximation simplifies the computation of the state posteriors.

In what follows, a bold case variable indicates the posterior expectation of the corresponding variable in italics. For example, 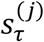 takes the value 0 or 1, while the posterior expectation 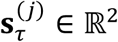 is the expected value of 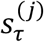 that lies between 0 and 1. Moreover, **a**^(*i,j*)^ ∈ ℝ^2×2^ denotes positive concentration parameters. Below, we use the posterior expectation of In *A*^(*i,j*)^ to encode posterior beliefs about the likelihood, which are given by

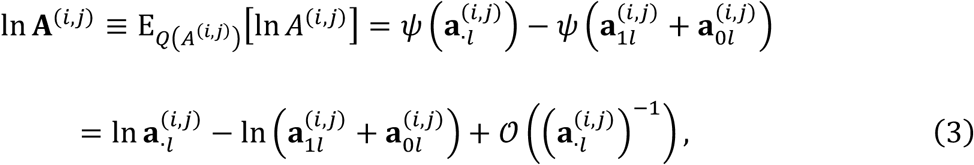

where *l* ∈ {1,0}. Here, *ψ*(·) ≡ Γ′(·)/Γ(·) denotes the digamma function, which arises naturally from the definition of the Dirichlet distribution. Please see (Friston et al., 2016) for details. 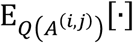 denotes the expectation over the posterior of *A*^(*i,j*)^.

The ensuing variational free energy of this generative model is then given by

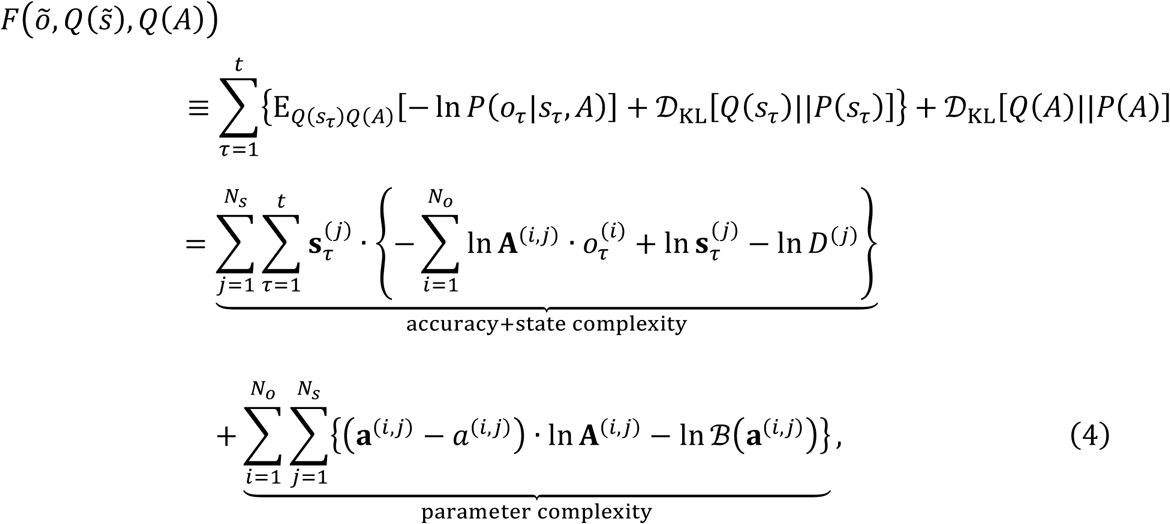

where 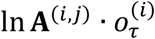 denotes the inner product of In **A**^(*i,j*)^ and a one-hot encoded vector of 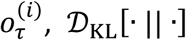 is the complexity as scored by the Kullback–Leibler divergence (Kullback & Leibler, 1951), and 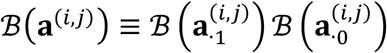 with 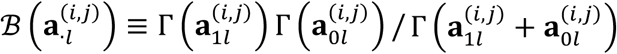 is the beta function. The first term in the final equality comprises the accuracy 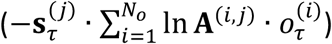 and (state) complexity 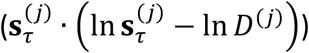. The accuracy term is simply the expected log likelihood of an observation, while complexity scores the divergence between prior and posterior beliefs. In other words, complexity reflects the degree of belief updating or degrees of freedom required to provide an accurate account of observations. Both belief updates to states and parameters incur a complexity cost: the state complexity increases with time *t*, while parameter complexity increases in the order of In *t*—and is thus negligible when *t* is large (see Supplementary Methods S1 for details). This means that we can ignore parameter complexity, when the scheme has experienced a sufficient number of outcomes. We will drop the parameter complexity in subsequent sections. In the remainder of this section, we show how the minimisation of variational free energy transforms (i.e., updates) priors into posteriors, when the parameter complexity is evaluated explicitly.

Inference optimises posterior expectations about the hidden states by minimising variational free energy. The optimal posterior expectations are obtained by solving the variation of *F* to give

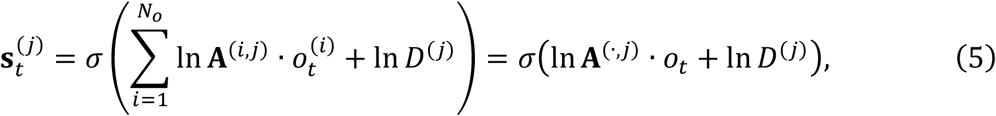

where *σ*(·) is the softmax function. As 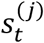 is a binary value in this work, the posterior expectation of 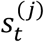 taking a value of one (ON state) can be expressed as

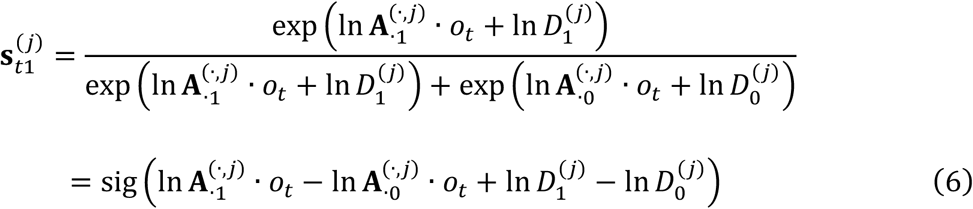

using the sigmoid function sig(z) ≡ 1/(1 + exp(−z)). Thus, the posterior expectation of 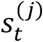 taking a value 0 (OFF state) is 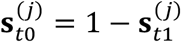. Here, 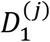 and 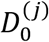 are constants denoting the prior beliefs about hidden states. Bayes optimal encoding is obtained when, and only when, the prior beliefs match the genuine prior distribution; i.e., 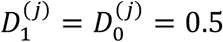 in this BSS setup. This concludes our treatment of inference about hidden states under this minimal scheme. Note that the updates in Equation (5) have a biological plausibility in the sense that the posterior expectations can be associated with nonnegative sigmoid-shape firing rates (also known as neurometric functions (Tolhurst et al., 1983; Newsome et al., 1989)), while the arguments of the sigmoid (softmax) function can be associated with neuronal depolarisation; rendering the softmax function a voltage-firing rate activation function. Please see (Friston, FitzGerald et al., 2017) for a more comprehensive discussion—and simulations using this kind of variational message passing to reproduce empirical phenomena; such as place fields, mismatch negativity responses, phase-precession, pre-play activity, etc in systems neuroscience.

In terms of learning, by solving the variation of *F* with respect to **a**^(*i,j*)^, the optimal posterior expectations about the parameters are given by

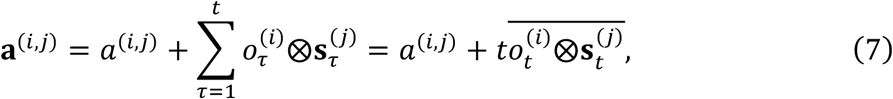

where *a*^(*i,j*)^ is the prior, 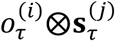 expresses the outer product of a one-hot encoded vector of 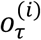 and 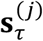, and 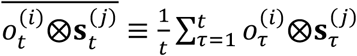. Thus, the optimal posterior expectation of matrix *A* is

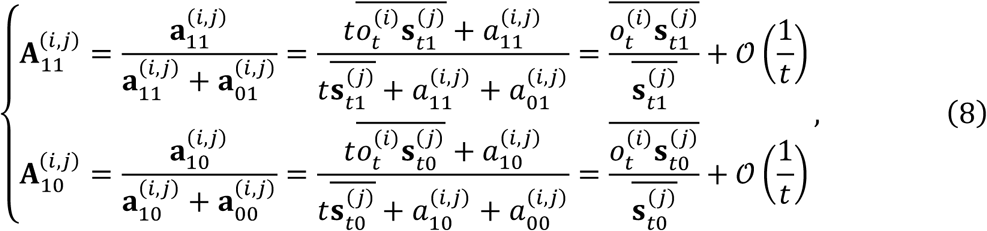

where 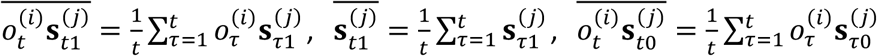, and 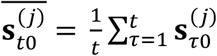. Further, 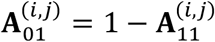 and 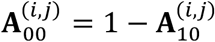. The prior of parameters *a*^(*i,j*)^ is in the order of 1 and is thus negligible when *t* is large. The matrix **A**^(*i,j*)^ express the optimal posterior expectations of 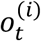 taking the ON state when 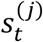 is ON 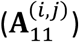 or OFF 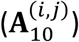, or 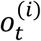 taking the OFF state when 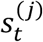 is ON 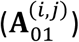 or OFF 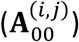. Although this expression may seem complicated, it is fairly straightforward. The posterior expectations of the likelihood simply accumulate posterior expectations about the co-occurrence of states and their outcomes. These accumulated (Dirichlet) parameters are then normalised to give a likelihood or probability. Crucially, one can observe the associative or Hebbian aspect of this belief update, expressed here in terms of the outer products between outcomes and posteriors about states in Equation (7). We now turn to the equivalent update for neural activities and synaptic weights of a neural network.

### 2.3 Neural activity and Hebbian plasticity models

Next, we consider the neural activity and synaptic plasticity in the neural network (Fig. 1B). We assume that the *j*-th neuron’s activity *x*_*tj*_ is given by

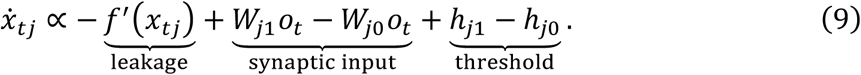

We suppose that 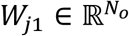 and 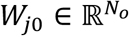 comprise row vectors of synapses, and *h*_*j*1_ ∈ ℝ and *h*_*j*0_ ∈ ℝ are adaptive thresholds that depend on the values of *W*_*j*1_ and *W*_*j*0_, respectively. One may regard *W*_*j*1_ and *W*_*j*0_ as excitatory and inhibitory synapses, respectively. We further assume that the nonlinear leakage *f*′(·) (i.e., the leak current) is the inverse of the sigmoid function (i.e., the logit function), such that the fixed point of *x*_*tj*_ is given by

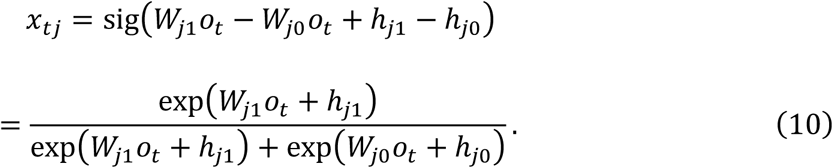

We further assume that synaptic strengths are updated following Hebbian plasticity with an activity-dependent homeostatic term as follows:

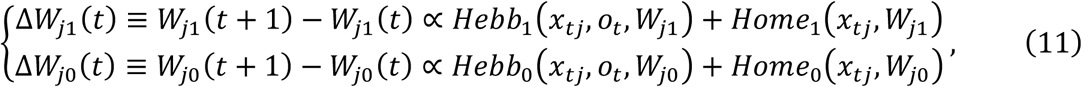

where *Hebb*_1_ and *Hebb*_0_ denote Hebbian plasticity as determined by the product of sensory inputs and neural outputs, and *Home*_1_ and *Home*_0_ denote homeostatic plasticity determined by output neural activity.

In the MDP scheme, posterior expectations about hidden states and parameters are usually associated with neural activity and synaptic strengths. Here, we can observe a formal similarity between the solutions for the state posterior (Equation (6)) and the activity in the neural network (Equation (10)). By this analogy, *x*_*tj*_ can be regarded as encoding the posterior expectation of the ON state 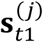. Moreover, *W*_*j*1_ and *W*_*j*0_ correspond to 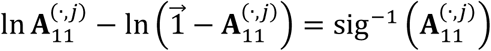 and 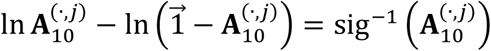, respectively, in the sense that they express the amplitude of *o*_*t*_ influencing *x*_*tj*_ or 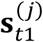. Here, 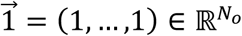 is a vector of ones. In particular, the optimal posterior of a hidden state taking a value of one (Equation (6)) is given by the ratio of the beliefs about ON and OFF states, expressed as a sigmoid function. Thus, to be a Bayes optimal encoder, the fixed point of neural activity needs to be a sigmoid function. This requirement is straightforwardly ensured when *f*′(*x*_*tj*_) is the inverse of the sigmoid function (see Equation (13) below). Under this condition, the fixed point or solution for *x*_*tk*_ (Equation (10)) compares inputs from ON and OFF pathways, and thus *x*_*tj*_ straightforwardly encodes the posterior of the *j*-th hidden state being ON (i.e., 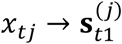). In short, the above neural network is effectively inferring the hidden state.

If the activity of the neural network is performing inference, does the Hebbian plasticity correspond to Bayes optimal learning? In other words, does the synaptic update rule in Equation (11) ensure that the neural activity and synaptic strengths asymptotically encode Bayes optimal posterior beliefs about hidden states 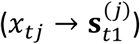 and parameters 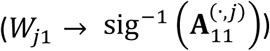, respectively? To this end, below we will identify a class of cost functions from which the neural activity and synaptic plasticity can be derived, and consider the conditions under which the cost function becomes consistent with variational free energy.

### 2.4 Neural network cost functions

Here, we consider a class of functions that constitute a cost function for both neural activity and synaptic plasticity. We start by assuming that the update of the *j*-th neuron’s activity (Equation (9)) is determined by the gradient of cost function *L*_*j*_; i.e., 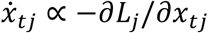. By integrating the right-hand side of Equation (9), we obtain a class of cost functions as

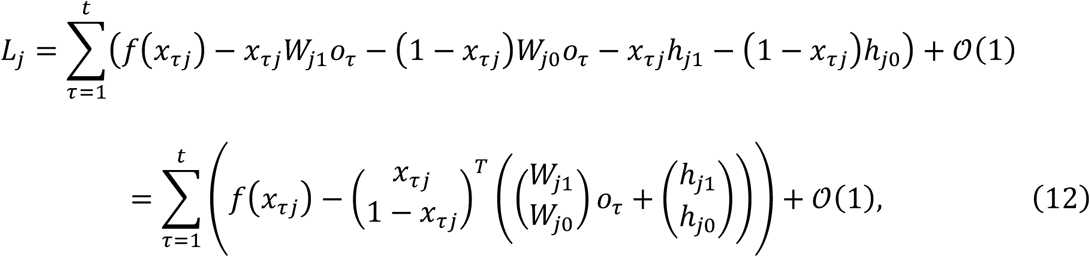

where the *𝒪*(1) term, which depends on *W*_*j*1_ and *W*_*j*0_, is of a lower order than the other terms (as they are *𝒪*(*t*)) and is thus negligible when *t* is large. Please see Supplementary Methods S3 for the case where we explicitly evaluate the *𝒪*(1) term, to demonstrate the formal correspondence between the initial values of synaptic strengths and the parameter prior *p*(*A*). The cost function of the entire network is defined by 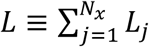. When *f*′(*x*_*τj*_) is the inverse of the sigmoid function, we have

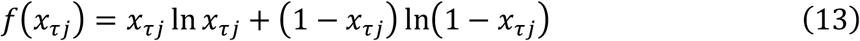

up to a constant term. We further assume that the synaptic weight update rule is derived from the same cost function *L*_*j*_. Thus, the synaptic plasticity is given by

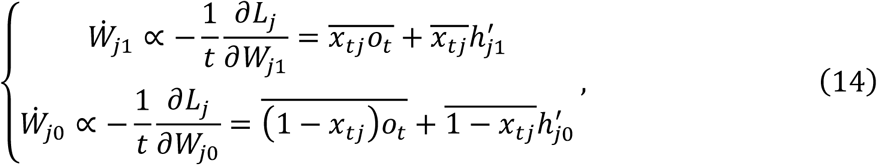

where 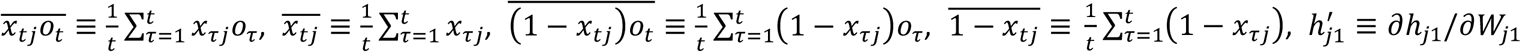, and 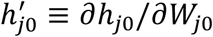. Note that the update of *W*_*j*1_ is not directly influenced by *W*_*j*0_, and *vice versa*, because they encode parameters in physically distinct pathways (i.e., the updates are local learning rules (Lee et al., 2000)). The update rule for *W*_*j*1_ can be viewed as Hebbian plasticity mediated by an additional activity-dependent term expressing homeostatic plasticity. Moreover, the update of *W*_*j*0_ can be viewed as anti-Hebbian plasticity with a homeostatic term, in the sense that *W*_*j*0_ is reduced when input (*o*_*t*_) and output (*x*_*tj*_) fire together. The fixed points of *W*_*j*1_ and *W*_*j*0_ are given by

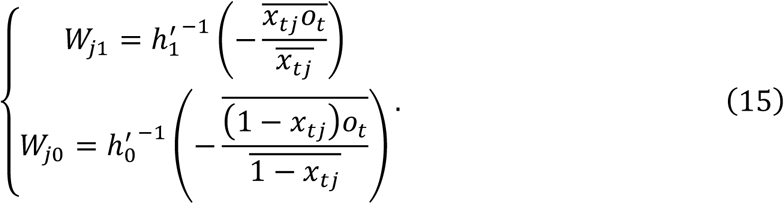

Crucially, these synaptic strength updates are a subclass of the general synaptic plasticity rule in Equation (11); see also Supplementary Methods S2 for the mathematical explanation. Therefore, if the synaptic update rule is derived from the cost function underlying neural activity, the synaptic update rule has a biologically plausible form comprising Hebbian plasticity and activity-dependent homeostatic plasticity.

### 2.5 Comparison with variational free energy

Here, we establish a formal relationship between the cost function *L* and variational free energy. We define 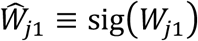 and 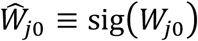 as the sigmoid functions of synaptic strengths. We consider the case in which neural activity is expressed as a sigmoid function and thus Equation (13) holds. As 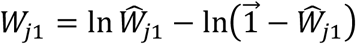, Equation (12) becomes

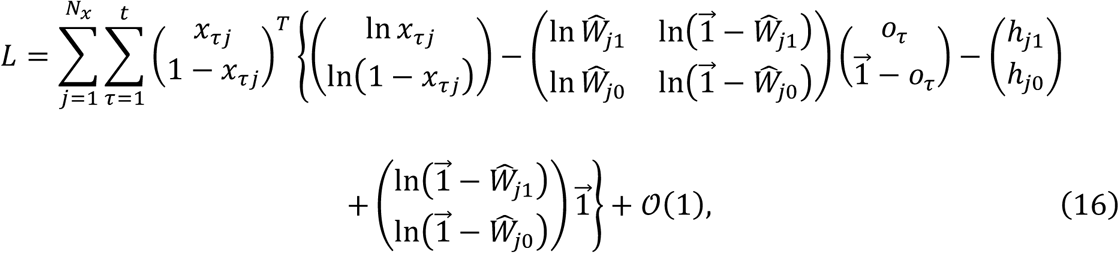

where 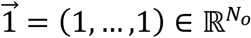. One can immediately see a formal correspondence between this cost function and variational free energy (Equation (4)). That is, when we assume that 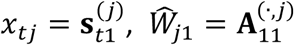, and 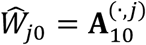, Equation (16) has exactly the same form as the sum of the accuracy and state complexity, which is the leading order term of variational free energy (see the first term in the last equality of Equation (4)).

Specifically, when the thresholds satisfy 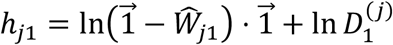 and 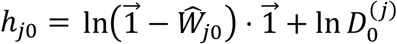, Equation (16) becomes equivalent to Equation (4) up to the In *t* order term (that disappears when *t* is large). Therefore, in this case, the fixed points of neural activity and synaptic strengths become the posteriors; thus, *x*_*tj*_ asymptotically becomes the Bayes optimal encoder for a large *t* limit (provided with *D* that matches the genuine prior *D*^*^).

In other words, we can define perturbation terms 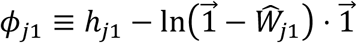 and 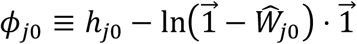 as functions of *W*_*j*1_ and *W*_*j*0_, respectively, and can express the cost function as

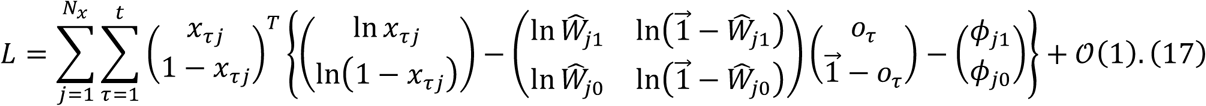

Here, without loss of generality, we can suppose that the constant terms in *ϕ*_*j*1_ and *ϕ*_*j*0_ are chosen to ensure that exp(*ϕ*_*j*1_) + exp(*ϕ*_*j*0_) = 1. Under this condition, (exp(*ϕ*_*j*1_), exp(*ϕ*_*j*0_)) can be viewed as the prior belief about hidden states

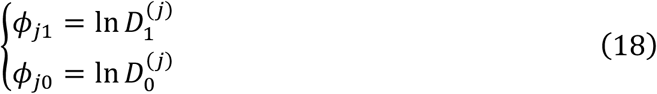

and thus Equation (17) is formally equivalent to the accuracy and state complexity terms of variational free energy.

This means that when the prior belief about states (*D*^(*j*)^) is a function of the parameter posteriors (**A**^(·,*j*)^), the generic cost function under consideration can be expressed in the form of variational free energy, up to the *𝒪*(ln *t*) term. A generic cost function *L* is sub-optimal from the perspective of Bayesian inference unless *ϕ*_*j*1_ and *ϕ*_*j*0_ are tuned appropriately to express the unbiased (i.e., optimal) prior belief. In this BSS setup, *ϕ*_*j*1_ = *ϕ*_*j*0_ = const is optimal; thus, a generic *L* would asymptotically give an upper bound of variational free energy with the optimal prior belief about states when *t* is large.

### 2.6 Analysis on synaptic update rules

To explicitly solve the fixed points of *W*_*j*1_ and *W*_*j*0_ that provide the global minimum of *L*, we suppose *ϕ*_*j*1_ and *ϕ*_*j*0_ as linear functions of *W*_*j*1_ and *W*_*j*0_, respectively, given by

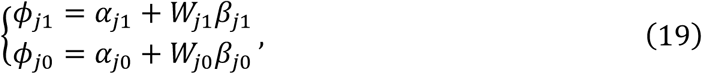

where *α*_*j*1_, *α*_*j*0_ ∈ ℝ and 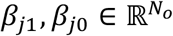 are constants. By solving the variation of *L* with respect to *W*_*j*1_ and *W*_*j*0_, we find the fixed point of synaptic strengths as

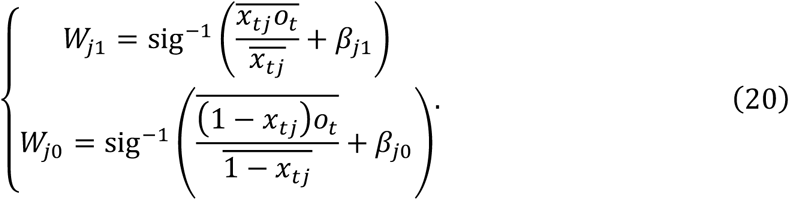

Since the update from *t* to *t*+1 is expressed as 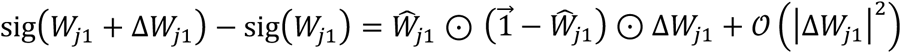 and 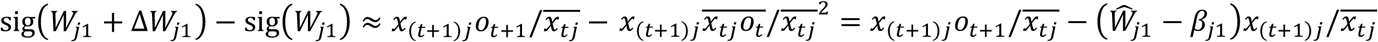, we recover the following synaptic plasticity:

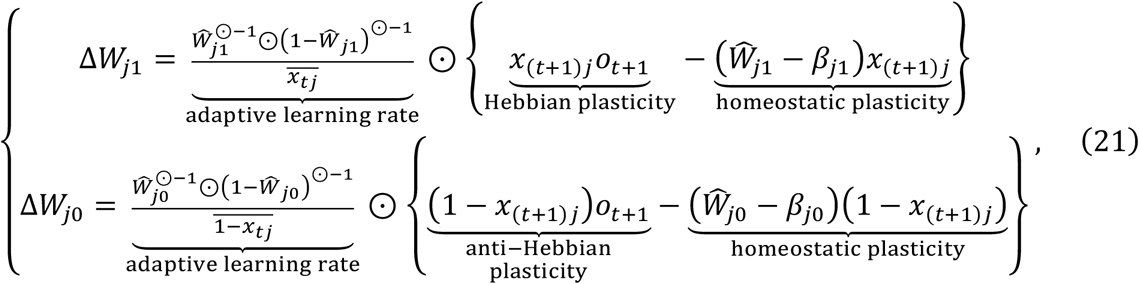

where ⊙ denotes the element-wise (Hadamard) product and 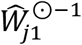 denotes the element-wise inverse of 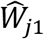. This synaptic plasticity rule is a subclass of the generic synaptic plasticity rule in Equation (11).

In summary, we demonstrated that under a few minimal assumptions and ignoring small contributions to weight updates, the neural network under consideration can be regarded as minimising an approximation to model evidence, because the cost function can be formulated in terms of variational free energy. In what follows, we will rehearse our analytic results and then use numerical analyses to illustrate Bayes optimal inference (and learning) in a neural network when, and only when, it has the right priors.

## 3. Results

### 3.1 Analytical form of neural network cost functions

The analysis in the preceding section rests on the following assumptions:

1. *Updates of neural activity and synaptic weights are determined by a gradient descent on a cost function L.*
2. *Neural activity is updated by the weighted sum of sensory inputs, and its fixed point is expressed as the sigmoid function.* Under these assumptions, we can express the cost function for a neural network as follows (see Equation (17)):

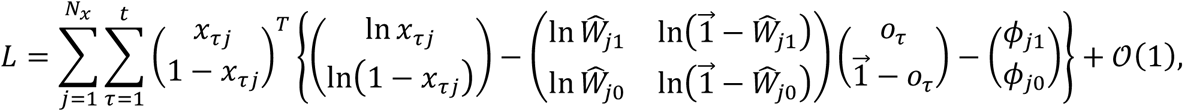

where 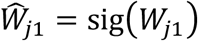 and 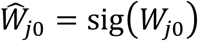 hold, and *ϕ*_*j*1_ and *ϕ*_*j*0_ are functions of *W*_*j*1_ and *W*_*j*0_, respectively. The log likelihood function (accuracy term) and divergence of hidden states (complexity term) of variational free energy emerge naturally under the assumption of a sigmoid activation function. The cost function above has additional terms denoted by *ϕ*_*j*1_ and *ϕ*_*j*0_. In other words, we can say that the cost function *L* is variational free energy under a sub-optimal prior belief about hidden states, depending on *W*_*j*1_ and 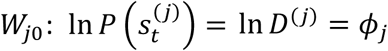, where *ϕ*_*j*_ ≡ (*ϕ*_*j*1_, *ϕ*_*j*0_). This prior alters the landscape of the cost function in a sub-optimal manner and thus provides a biased solution for neural activities and synaptic strengths, which differ from the Bayes optimal encoders. For analytical tractability, we further assume the following:
3. *The perturbation terms* (*ϕ*_*j*1_ *and ϕ*_*j*0_) *that constitute the difference between the cost function and variational free energy with optimal prior beliefs can be expressed as linear equations of W*_*j*1_ *and W*_*j*0_. From assumption 3, Equation (17) becomes

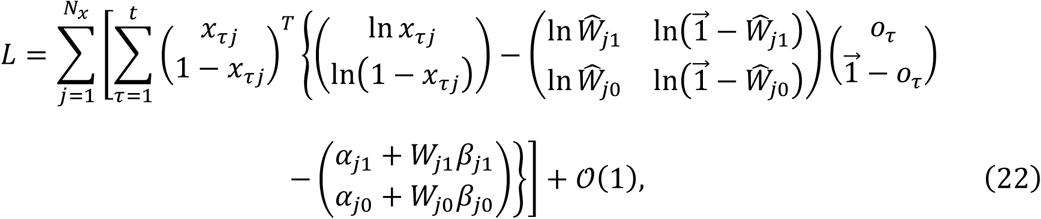

where {*α*_*j*1,_ *α*_*j*0,_ *β*_*j*1,_ *β*_*j*0_} are constants. The cost function has degrees of freedom with respect to the choice of constants {*α*_*j*1,_ *α*_*j*0,_ *β*_*j*1,_ *β*_*j*0_}, which correspond to the prior belief about states *D*^(*j*)^. The neural activity and synaptic strengths that give the minimum of a generic physiological cost function *L* are biased by these constants, which may be analogous to physiological constraints (see Discussion for details).

The cost function of the neural networks considered is characterised only by *ϕ*_*j*_. Thus, after fixing *ϕ*_*j*_ by fixing constrains (*α*_*j*1,_ *α*_*j*0_) and (*β*_*j*1,_ *β*_*j*0_), the remaining degrees of freedom are the initial synaptic weights. These correspond to the prior distribution of parameters *P*(*A*) in the variational Bayesian formulation (please see Supplementary Methods 3).

The fixed point of synaptic strengths that give the minimum of *L* is given analytically as Equation (20), expressing that (*β*_*j*1,_ *β*_*j*0_) deviates the centre of the nonlinear mapping—from Hebbian products to synaptic strengths—from the optimal position (shown in Equation (8)). As shown in Equation (14), the derivative of *L* with respect to *W*_*j*1_ and *W*_*j*0_ recovers the synaptic update rules that comprise Hebbian and activity-dependent homeostatic terms. Although Equation (14) expresses the dynamics of synaptic strengths that converge to the fixed point, it is consistent with a plasticity rule that gives the synaptic change from *t* to *t*+1 (Equation (21)).

Hence, based on assumptions 1 and 2, we find that the cost function approximates variational free energy; see also Supplementary Table S1 for their correspondence. Under this condition, neural activity encodes the posterior expectation about hidden states, 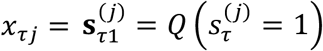, and synaptic strengths encode the posterior expectation of the parameters, 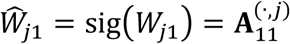 and 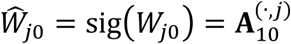. In addition, based on assumption 3, the accuracy of approximation depends on the deviation of constants {*α*_*j*1,_ *α*_*j*0,_ *β*_*j*1,_ *β*_*j*0_} from their optimal values. From a Bayesian perspective, these constants can be viewed as prior beliefs, 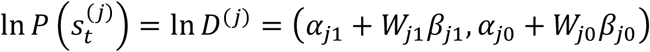, when we assume that (*x*_*tj*_, 1 − *x*_*tj*_) represents the state posterior 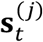. When and only when (*α*_*j*1,_ *α*_*j*0_) = (− ln 2, − ln 2) and 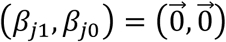, the cost function becomes variational free energy with optimal prior beliefs (for BSS), whose global minimum ensures Bayes optimal encoding.

In short, we identify a class of biologically plausible cost functions from which the update rules for both neural activity and synaptic plasticity can be derived. When the activation function for neural activity is a sigmoid function, a cost function in this class is expressed straightforwardly as variational free energy. With respect to the choice of constants expressing physiological constraints in the neural network, the cost function has degrees of freedom that may be viewed as (potentially sub-optimal) prior beliefs from the Bayesian perspective. Now, we illustrate the implicit inference and learning in neural networks through simulations of BSS.

### 3.2 Numerical simulations

Here, we simulated the dynamics of neural activity and synaptic strengths when they followed a gradient descent on the cost function in Equation (22). We considered a BSS comprising two hidden sources (or states) and 32 observations (or sensory inputs), formulated as an MDP. The two hidden sources comprised four patterns: 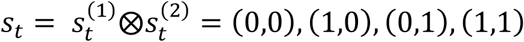. An observation 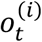 was generated through the likelihood mapping *A*^(*i*)^, defined as

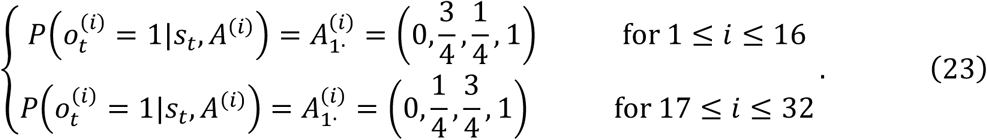

Here, for example, 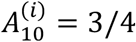 for 1 ≤ *i* ≤ 16 is the probability of 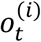 taking one when *s*_*t*_ = (1,0). The simulations continued over *T* = 10^4^ time steps. Notably, this simulation setup is exactly the same experimental setup as that we used for *in vitro* neural networks (Isomura et al., 2015; Isomura, Friston, 2018). We leverage this setup to clarify the relationship among our empirical work, a feed-forward neural network model, and variational Bayesian formulations.

First, as in (Isomura & Friston, 2018), we demonstrated that a network with a cost function with optimised constants ((*α*_*j*1,_ *α*_*j*0_) = (− ln 2, − ln 2) and 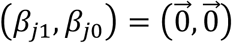) can perform BSS successfully (Fig. 2). The responses of neuron 1 came to recognise source 1 after training, indicating that neuron 1 learnt to encode source 1 (Fig. 2A). Meanwhile, neuron 2 learnt to infer source 2 (Fig. 2B). This demonstrates that minimisation of the cost function, with optimal constants, is equivalent to variational free energy minimisation, and hence is sufficient to emulate BSS. Next, we quantified the dependency of BSS performance on the form of the cost function, by varying the above-mentioned constants (Fig. 3).

**Figure 2.**
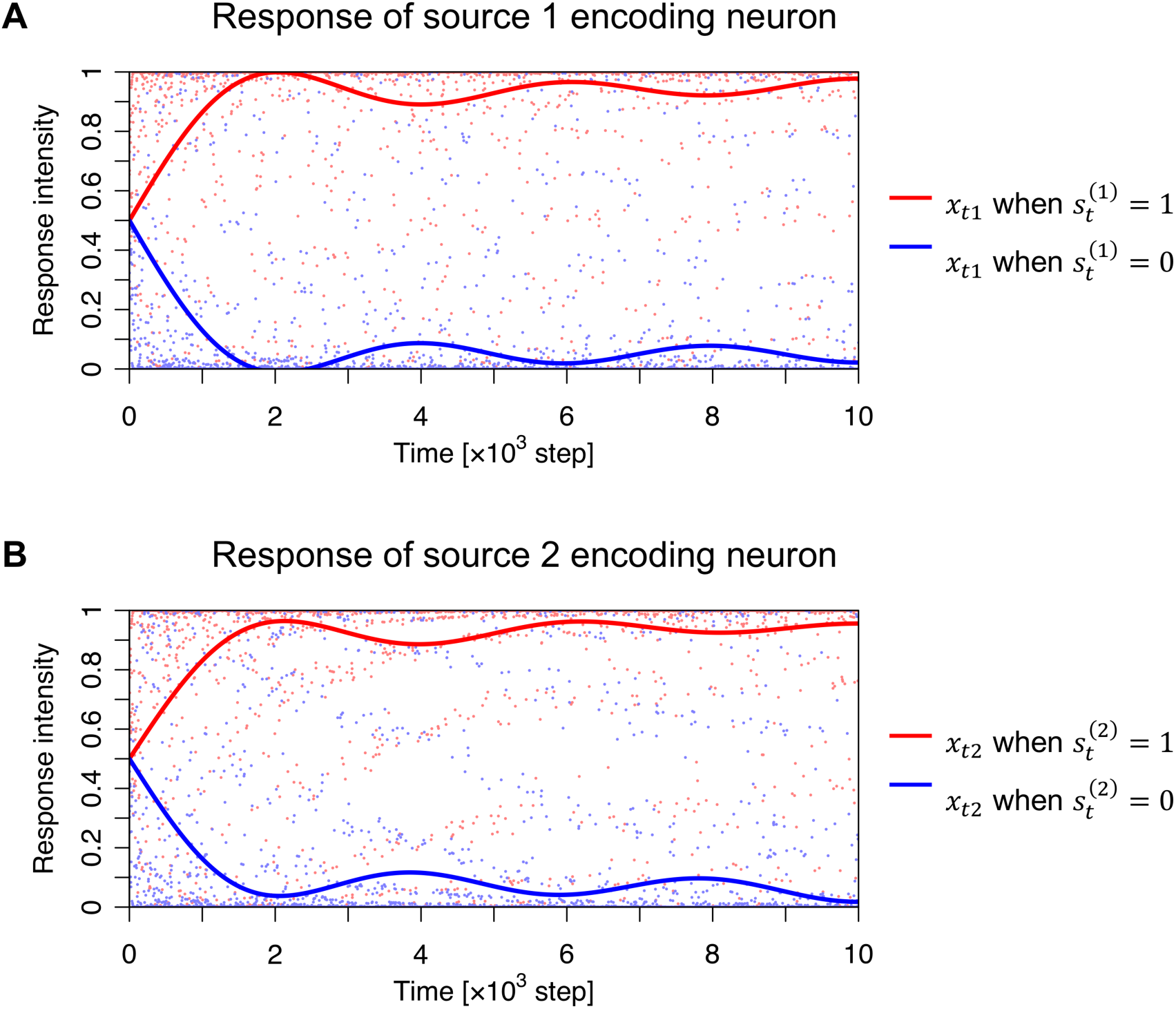
Emergence of response selectivity for a source. (**A**) Evolution of neuron 1’s responses that learn to encode source 1, in the sense that the response is high when source 1 takes a value of one (red dots), and it is low when source 1 takes a value of zero (blue dots). Lines correspond to smoothed trajectories obtained using a discrete cosine transform. (**B**) Emergence of neuron 2’s response that learns to encode source 2. These results indicate that the neural network succeeded in separating two independent sources. The code is provided as Supplementary Source Code.

**Figure 3.**
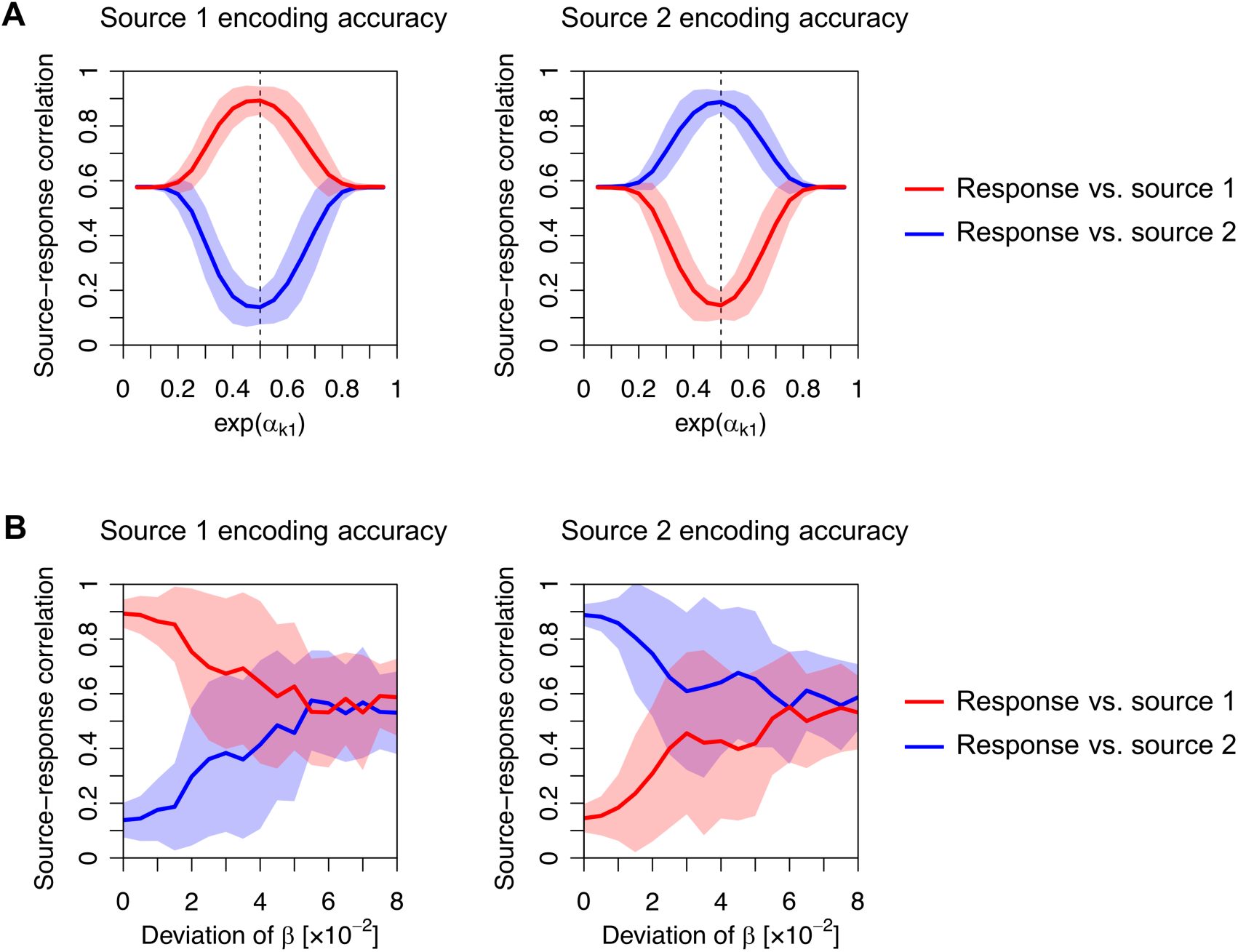
Dependence of source encoding accuracy on constants. Left panels show the magnitudes of the correlations between sources and responses of a neuron expected to encode source 1: 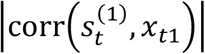 and 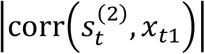. The right panels show the magnitudes of the correlations between sources and responses of a neuron expected to encode source 2: 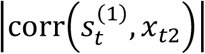 and 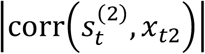. (**A**) Dependence on the constant *α* that controls the excitability of a neuron, when *β* is fixed to zero. The dashed line (0.5) indicates the optimal value of exp(*α*_*j*1_). (**B**) Dependence on constant *β*, when *α* is fixed as (*α*_*j*1,_ *α*_*j*0_) = (− ln 2, − ln 2). Elements of *β* were randomly generated from a Gaussian distribution with zero mean. The standard deviation of *β* was varied (horizontal axis), where zero deviation was optimal. Lines and shaded areas indicate the mean and standard deviation of the source-response correlation, evaluated with 50 different sequences. The code is provided as Supplementary Source Code.

We varied (*α*_*j*1,_ *α*_*j*0_) in a range of 0.05 ≤ exp(*α*_*j*1_) ≤ 0.95, while maintaining exp(*α*_*j*1_) + exp(*α*_*j*0_) = 1, and found that changing (*α*_*j*1,_ *α*_*j*0_) from (− ln 2, − ln 2) led to a failure of BSS. Because neuron 1 encodes source 1 with optimal *α*, the correlation between source 1 and the response of neuron 1 is close to one, while the correlation between source 2 and the response of neuron 1 is nearly zero. In the case of sub-optimal *α*, these correlations fall to around 0.5, indicating that the response of neuron 1 encodes a mixture of source 1 and source 2 (Fig. 3A). Moreover, a failure of BSS can be induced when the elements of *β* take values far from zero (Fig. 3B). When the elements of *β* are generated from a zero-mean Gaussian distribution, the accuracy of BSS—measured using the correlation between sources and responses—decreases as the standard deviation increases.

Our numerical analysis, under assumptions 1–3 mentioned above, shows that a network needs to employ a cost function that entails optimal prior beliefs to perform BSS, or equivalently, causal inference. Such a cost function is obtained when its constants, which do not appear in the variational free energy with the optimal generative model for BSS, become negligible. The important message here is that, in this setup, a cost function equivalent to variational free energy is necessary for Bayes optimal inference (Friston et al., 2006; Friston, 2010).

### 3.3 Phenotyping networks

We have shown that variational free energy (under the MDP scheme) is within the class of biologically plausible cost functions found in neural networks. The neural network’s parameters *ϕ*_*j*_ = ln *D*^(*j*)^ determine how the synaptic strengths change depending on the history of sensory inputs and neural outputs; thus, the choice of *ϕ*_*j*_ provides degrees of freedom in the shape of the generic cost functions under consideration that determine the purpose or function of the neural network. Among various *ϕ*_*j*_, only *ϕ*_*j*_ = (− ln 2, − ln 2) can make the cost function variational free energy with optimal prior beliefs for BSS. Hence, one could regard generic neural networks (of the sort considered in this paper) as performing approximate Bayesian inference under priors that may or may not be optimal. This result is as predicted by the complete class theorem as it implies that any response of a neural network is Bayes optimal under some prior beliefs (and cost function). Therefore, under the theorem, in principle, any neural network of this kind is optimal, when its prior beliefs are consistent with the process that generates outcomes. This perspective indicates the possibility of characterising a neural network model—and indeed a real neuronal network—in terms of its implicit prior beliefs.

These considerations raise the possibility of using empirically observed neuronal responses to infer the prior beliefs implicit in a neuronal network. For example, the synaptic matrix (*W*_*j*1_, *W*_*j*0_) can be estimated statistically from response data. By plotting its trajectory over the training period as a function of the history of a Hebbian product, one can estimate the cost function constants. If these constants express a near-optimal *ϕ*_*j*_, it can be concluded that the network has, effectively, the right sort of priors for BSS. As we have shown analytically and numerically, a cost function with (*α*_*j*1,_ *α*_*j*0_) far from (− ln 2, − ln 2) or a large deviation of (*β*_*j*1,_ *β*_*j*0_) does not provide the Bayes optimal encoder for performing BSS. Since actual neuronal networks can perform BSS (Isomura et al., 2015; Isomura & Friston, 2018), it can be envisaged that the implicit cost function will exhibit a near-optimal *ϕ*_*j*_.

One can pursue this analysis further and model the responses or decisions of a neural network using the above-mentioned Bayes optimal MDP scheme under different priors. Thus, the priors in the MDP scheme can be adjusted to maximise the likelihood of empirical responses. This sort of approach has been used in system neuroscience to characterise the choice behaviour in terms of subject specific priors. Please refer to (Schwartenbeck & Friston, 2016) for further details.

Finally, from a practical perspective for optimising neural networks, understanding the formal relationship between cost functions and variational free energy enables us to specify the optimum value of any free parameter to realize some functions. In the present setting, we can effectively optimise the constants by updating the priors themselves, such that they minimise the variational free energy for BSS. Under the Dirichlet form for the priors, the implicit threshold constants of the objective function can then be optimised using the following updates:

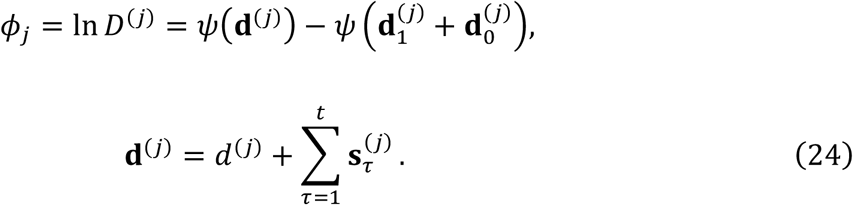

Please refer to (Schwartenbeck & Friston, 2016) for further details. In effect, this update will simply add the Dirichlet concentration parameters, 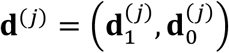, to the priors in proportion to the temporal summation of the posterior expectations about the hidden states. Therefore, by committing to cost functions that underlie variational inference and learning, any free parameter can be updated in a Bayes optimal fashion when a suitable generative model is available.

## 4. Discussion

In this work, we investigated a class of biologically plausible cost functions for neural networks. A single-layer feed-forward neural network with a sigmoid activation function that receives sensory inputs generated by hidden states (i.e., BSS setup) was considered. We identified a class of cost functions by assuming that neural activity and synaptic plasticity minimise a common function *L*. The derivative of *L* with respect to synaptic strengths furnishes a synaptic update rule following Hebbian plasticity, equipped with activity-dependent homeostatic term. We have shown that the dynamics of a single-layer feed-forward neural network—that minimises its cost function—is asymptotically equivalent to that of variational Bayesian inference under a particular but generic (latent variable) generative model. Hence, the cost function of the neural network can be viewed as variational free energy, and biological constraints that characterise the neural network—in the form of thresholds and neuronal excitability—become prior beliefs about hidden states. This relationship holds regardless of the true generative process of the external world. We have focused on discrete latent variable models that can be regarded as special (reduced) cases of partially observable Markov decision processes (POMDP). However, because our treatment is predicated on the complete class theorem (Brown, 1981; Wald, 1947), the same conclusions should, in principle, be reached when using continuous state space models. Within the class of discrete state space models, it is fairly straightforward to generate continuous outcomes from discrete latent states; as exemplified by discrete variational autoencoders (Rolfe, 2016) or mixed models, as described in (Friston, Parr et al., 2017).

One can understand the nature of the constants {*α*_*j*1,_ *α*_*j*0,_ *β*_*j*1,_ *β*_*j*0_} from the biological and Bayesian perspectives as follows: (*α*_*j*1,_ *α*_*j*0_) determines the firing threshold and thus controls the mean firing rates. In other words, these parameters control the amplitude of excitatory and inhibitory inputs, which may be analogous to the roles of GABAergic inputs and neuromodulators in biological neuronal networks (Pawlak et al., 2010; Frémaux & Gerstner, 2016; Kuśmierz et al., 2017). At the same time, (*α*_*j*1,_ *α*_*j*0_) encodes prior beliefs about states, which exert a large influence on the state posterior. The state posterior is biased if (*α*_*j*1,_ *α*_*j*0_) is selected in a sub-optimal manner—in relation to the process that generates inputs. Meanwhile, (*β*_*j*1,_ *β*_*j*0_) determines the accuracy of synaptic strengths that represent the likelihood mapping of an observation 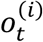 taking 1 (ON state) depending on hidden states (please compare Equation (8) and Equation (20)). Under a usual MDP setup where the state prior does not depend on the parameter posterior, the encoder becomes Bayes optimal when and only when 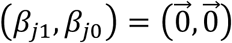. These constants can represent biological constraints on synaptic strengths, such as the range of spine growth, spinal fluctuations, or the effect of synaptic plasticity induced by spontaneous activity independent of external inputs. Although the fidelity of each synapse is limited due to such internal fluctuations, the accumulation of information over a large number of synapses should allow accurate encoding of hidden states in the current formulation.

In previous reports, we have shown that *in vitro* neural networks—comprising a cortical cell culture—perform BSS when receiving electrical stimulations generated from two hidden sources (Isomura et al., 2015). Furthermore, we showed that minimising variational free energy under an MDP is sufficient to reproduce the learning observed in an *in vitro* network (Isomura & Friston, 2018). Our framework for identifying biologically plausible cost functions could be relevant for identifying the principles that underlie learning or adaptation processes in biological neuronal networks, using empirical response data. Here, we illustrated this potential in terms of the choice of function *ϕ*_*j*_ in the cost functions *L*. In particular, if *ϕ*_*j*_ is close to a constant (− ln 2, − ln 2), the cost function is expressed straightforwardly as a variational free energy with small state prior biases. In the future work, we plan to apply this scheme to empirical data and examine the biological plausibility of variational free energy minimisation.

The correspondence highlighted in this work enables one to identify a generative model (comprising likelihood and priors) that a neural network is using. The formal correspondence between neural network and variational Bayesian formations rests on the asymptotic equivalence between the neural network’s cost functions and variational free energy (under some priors). Although variational free energy can take an arbitrary form, the correspondence provides biologically plausible constraints for neural networks that implicitly encode prior distributions. Hence, this formulation is potentially useful for identifying the implicit generative models that underlie the dynamics of real neuronal circuits. In other words, one can quantify the dynamics and plasticity of a neuronal circuit in terms of variational Bayesian inference and learning under an implicit generative model.

The dependence between the likelihood function and the state prior vanishes when the network uses the optimal threshold to perform inference with a generative process that does not involve dependence between the likelihood and the state prior. In other words, the dependence arises from the sub-optimality of the choice of the state prior. This means that the dependence is due to the degrees of freedom in the choice of the threshold that a neural network and its cost function possess. Nevertheless, minimisation of the cost function can render the network Bayes optimal in the variational Bayesian sense, including the choice of the state prior, as described in the previous section. This is because only variational free energy with the optimal priors provides the minimum among a class of neural network cost functions under consideration.

Although we have described the generative process in terms of an MDP, we have ignored state transitions. This means the generative model in this paper reduces to a simple latent variable model, with categorical states and outcomes. As noted above, we refer to MDP models because they predominate in descriptions of variational (Bayesian) belief updating; e.g., (Friston, FitzGerald et al., 2017). Clearly, many generative processes entail state transitions, leading to hidden Markov models (HMM). When state transitions depend upon control variables, we have a POMDP. To deal with such cases, extensions of the current framework are required, which we hope to consider in future work.

In summary, we first identified a class of biologically plausible cost functions for neural networks that underlie changes in both neural activity and synaptic plasticity. We then identified an asymptotic equivalence between these cost functions and the cost functions used in variational Bayesian formations. Given this equivalence, changes in the activity and synaptic strengths of a neuronal network can be viewed as Bayesian belief updating; namely, a process of transforming priors over hidden states and parameters into posteriors, respectively. Hence, a cost function in this class becomes Bayes optimal when activity thresholds correspond to appropriate priors in an implicit generative model. In short, the neural and synaptic dynamics of neural networks can be cast as inference and learning, under a variational Bayesian formation. This is potentially important for two reasons. First, it means that there are some threshold parameters for any neural network (in the class considered) that can be optimised for applications to data, when there are precise prior beliefs about the process generating those data. Second, in virtue of the complete class theorem, one can reverse engineer the priors that any neural network is adopting. This may be interesting when real neuronal networks can be modelled using neural networks of the class that we have considered. In other words, if one can fit neuronal responses—using a neural network model parameterised in terms of threshold constants—it becomes possible to evaluate the implicit priors using the above equivalence. This may find a useful application when applied to *in vitro* (or *in vivo*) neuronal networks (Isomura, Friston, 2018; Levin, 2013) or, indeed, dynamic causal modelling of distributed neuronal responses from non-invasive data (Daunizeau et al., 2011). In this context, the neural network can, in principle, be used as a dynamic causal model to estimate threshold constants and implicit priors. This ‘reverse engineering’ speaks to estimating the priors used by real neuronal systems, under ideal Bayesian assumptions; sometimes referred to as meta Bayesian inference (Daunizeau et al., 2010).

## Acknowledgements

T.I. is funded by RIKEN Center for Brain Science. K.J.F. is funded by a Wellcome Principal Research Fellowship (Ref: 088130/Z/09/Z). The funders had no role in study design, data collection and analysis, decision to publish, or preparation of the manuscript.

## Supplementary Information

### Supplementary Tables

**Table S1.** Correspondence of variables and functions.

| Neural network formation |  | Variational Bayes formation |  |
| --- | --- | --- | --- |
| Neural activity | $x_{tj} \Leftrightarrow$ | $\mathbf{s}_{t1}^{(j)}$ | State posterior |
| Sensory input | $o_t \Leftrightarrow$ | $o_t$ | Observation |
| Synaptic strength | $W_{j1} \Leftrightarrow$ | $\text{sig}^{-1}(\mathbf{A}_{11}^{(\cdot,j)})$ | |
| | $\widehat{W}_{j1} \Leftrightarrow$ | $\mathbf{A}_{11}^{(\cdot,j)}$ | Parameter posterior |
| Perturbation term | $\phi_{j1} \Leftrightarrow$ | $\ln D_1^{(j)}$ | State prior |
| Threshold | $h_{j1} \Leftrightarrow$ | $\ln(\vec{1} - \mathbf{A}_{11}^{(\cdot,j)}) \cdot \vec{1} + \ln D_1^{(j)}$ | |
| Initial synaptic strengths | $\lambda_{j1} \odot \widehat{W}_{j1}^{init} \Leftrightarrow$ | $a_{11}^{(\cdot,j)}$ | Parameter prior |

### Supplementary Methods

#### S1. Order of the parameter complexity

The order of the parameter complexity term

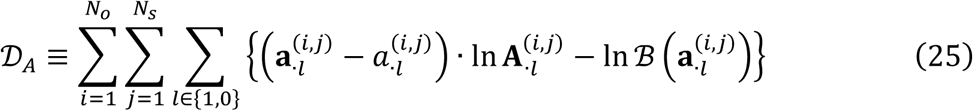

is computed. To avoid the divergence of In 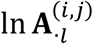, all the elements of 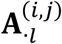 are assumed to be larger than a positive constant *ε*. This means that all the elements of 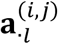 are in the order of *t*. The first term of Equation (25) becomes 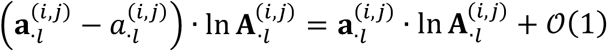 since 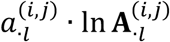 is in the order of 1. Moreover, from Equation (3), 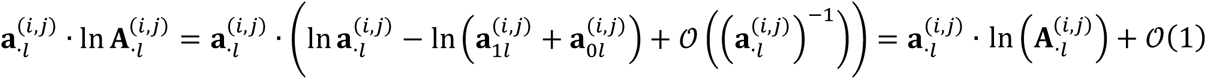. Meanwhile, the second term of Equation (25) comprises the logarithms of gamma functions as In 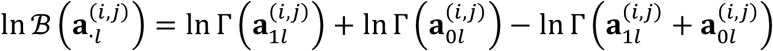. From Stirling’s formula,

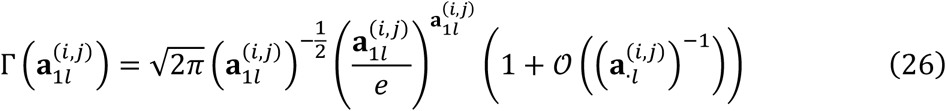

holds. The logarithm of 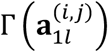 is evaluated as

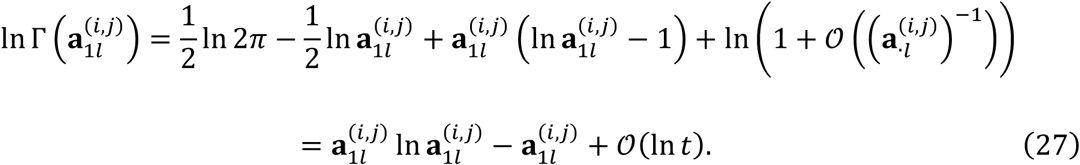

Similarly, 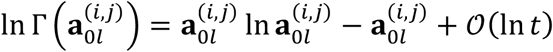 and 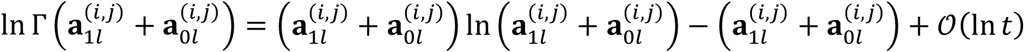 hold. Thus, we obtain

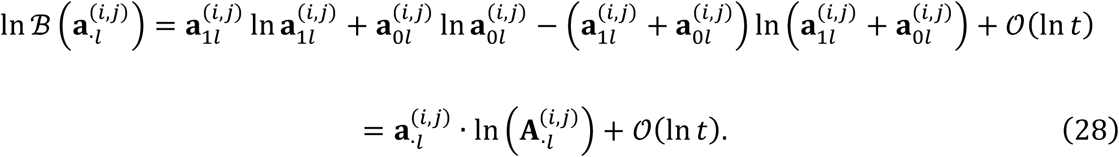

Hence, Equation (25) becomes

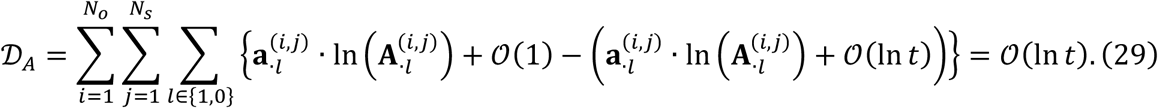

Therefore, we obtain

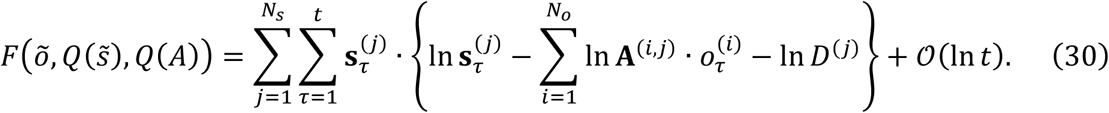

Under the current generative model comprising binary hidden states and binary observations, the optimal posterior expectation of **A** can be obtained up to the order of ln *t* /*t* even when the 𝒪(ln *t*) term in Equation (30) is ignored. Solving the variation of *F* with respect to 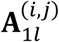 yields the optimal posterior expectation. From 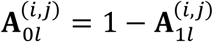, we find

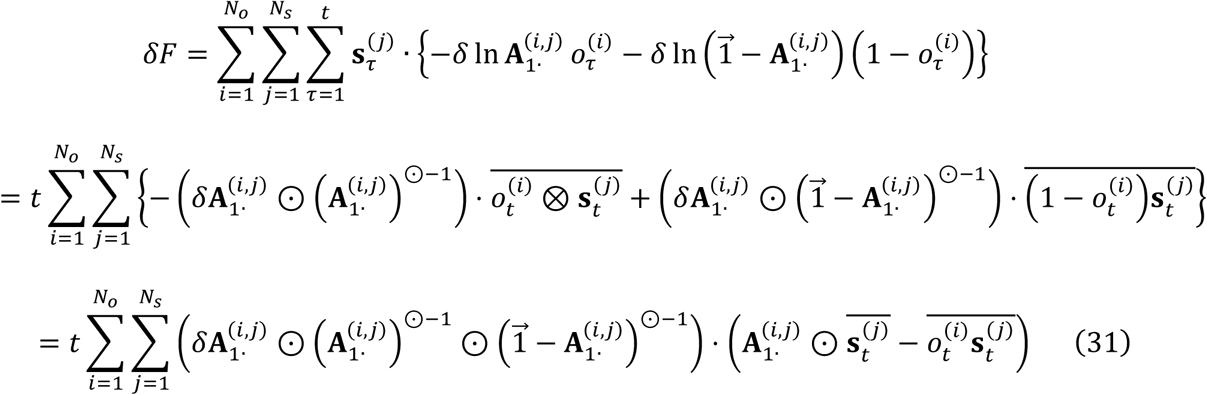

up to the order of ln *t*. Here, 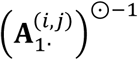 denotes the element-wise inverse of 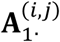. From *δF* = 0, we find

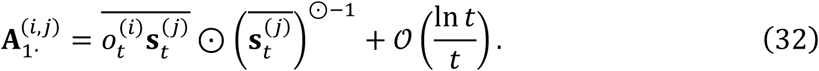

Therefore, we obtain the same result as Equation (8) up to the order of ln *t* /*t*.

#### S2. Derivation of synaptic plasticity rule

We consider synaptic strengths at time *t, W*_*j*1_ = *W*_*j*1_(*t*), and define the change as Δ*W*_*j*1_ ≡ *W*_*j*1_(*t* + 1) − *W*_*j*1_(*t*). From Equation (15), 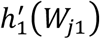 satisfies both

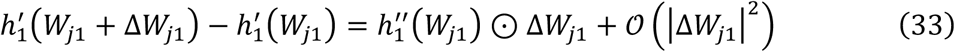

and

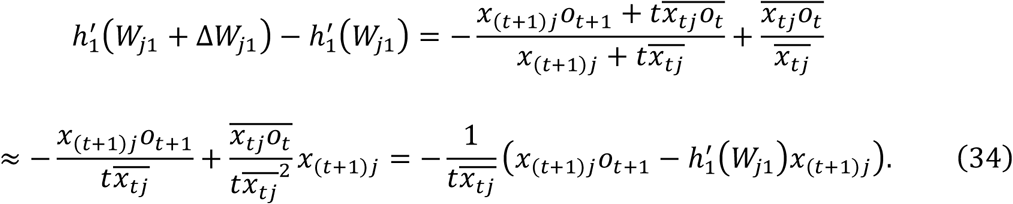

Thus, we find

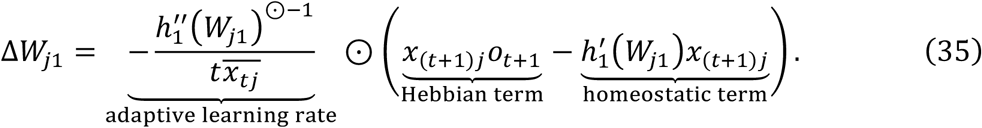

Similarly,

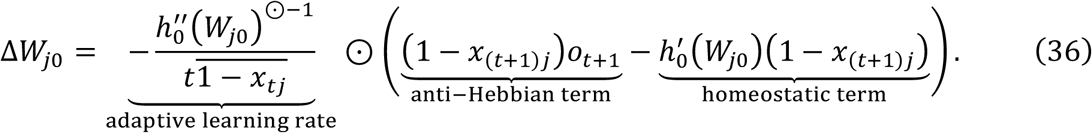

These plasticity rules express (anti-) Hebbian plasticity with a homeostatic term.

#### S3. Correspondence between parameter prior distribution and initial synaptic strengths

In general, optimising a model of observable quantities—including a neural network—can be cast inference, if there exists a learning mechanism that updates the hidden states and parameters of that model based on observations. (Exact and variational) Bayesian inference treats the hidden states and parameters as random variables, and thus transforms prior distributions *P*(*S*_*t*_), *P*(*A*) into posteriors *Q*(*S*_*t*_), *Q*(*A*). In other words, Bayesian inference is a process of transforming the prior to the posterior based on observations *o*_1_, …, *o*_*t*_ under a generative model. From this perspective, the incorporation of prior knowledge about the hidden states and parameters is an important aspect of Bayesian inference.

The minimisation of a cost function by a neural network updates its activity and synaptic strengths based on observations under the given network properties (e.g., activation function and thresholds). According to the complete class theorem, this process can always be viewed as Bayesian inference. In the main text, we demonstrated that a class of cost functions—for a single-layer feed-forward network with a sigmoid activation function—has a form equivalent to variational free energy under a particular latent variable model. Here, neural activity *x*_*t*_ and synaptic strengths *W* come to encode the posterior distributions over hidden states *Q*′(*S*_*t*_) and parameters *Q*′(*A*), respectively, where *Q*′(*S*_*t*_) and *Q*′(*A*) follow categorical and Dirichlet distributions, respectively. Moreover, we identified that the perturbation factors *ϕ*_*j*_ —that characterise the threshold function—correspond to the logarithm of the state prior *P*(*S*_*t*_) expressed as a categorical distribution.

However, one might ask whether the posteriors obtained using the network *Q*′(*S*_*t*_), *Q*′(*A*) are formally different from those obtained using variational Bayesian inference *Q*(*S*_*t*_), *Q*(*A*), since only the latter explicitly considers the prior distribution of parameters *P*(*A*). Thus, one may wonder if the network merely influences update rules that are similar to variational Bayes but does not transform the priors *P*(*S*_*t*_), *P*(*A*) into the posteriors *Q*(*S*_*t*_), *Q*(*A*), despite the asymptotic equivalence of the cost functions.

Below, we show that the initial values of synaptic strengths 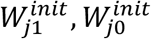 correspond to the parameter prior *P*(*A*) expressed as a Dirichlet distribution, to show that a neural network indeed transforms the priors into the posteriors. For this purpose, we specify the order 1 term in Equation (12) to make the dependence on the initial synaptic strengths explicit. Specifically, we modify Equation (12) as

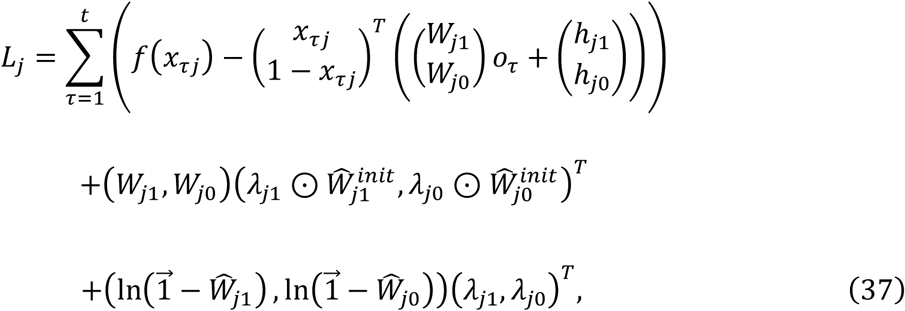

where 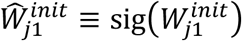 and 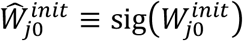 are the sigmoid functions of the initial synaptic strengths, and 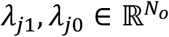 are row vectors of the inverse learning rate factors that express the insensitivity of the synaptic strengths to the activity-dependent synaptic plasticity. The third term of Equation (37) expresses the integral of 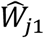 and 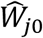 (with respect to *W*_*j*1_ and *W*_*j*0_, respectively). This ensures that when *t* = 0 (i.e., when the first term on the right-hand side of Equation (37) is zero), the derivative of *L*_*j*_ is given by 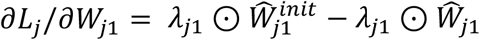, and thus 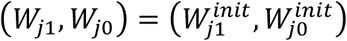 provides the fixed point of *L*_*j*_.

Similar to the transformation from Equation (12) to Equation (17), we compute Equation (37) as

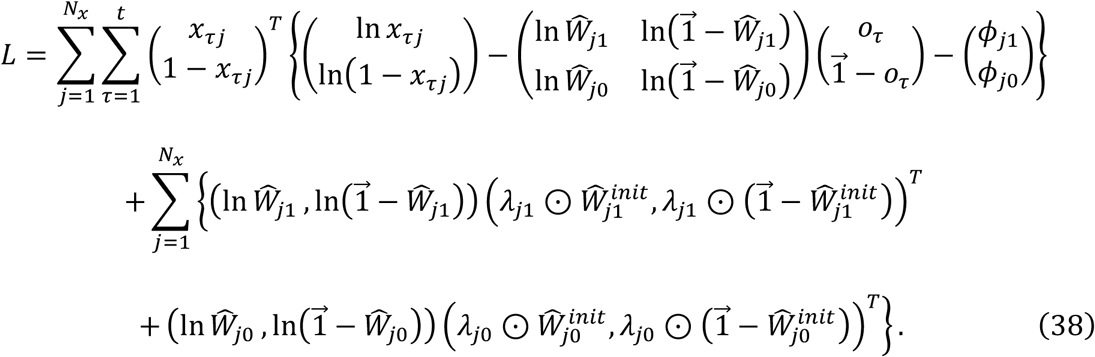

Note that we used 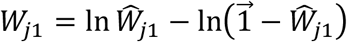. Crucially, analogous to the correspondence between 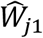 and the Dirichlet parameters of the parameter posterior 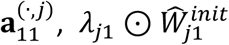 can be formally associated with the Dirichlet parameters of the parameter prior 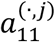. Hence, one can see the formal correspondence between the second and third terms on the right-hand side of Equation (38) and the expectation of the log parameter prior in Equation (4):

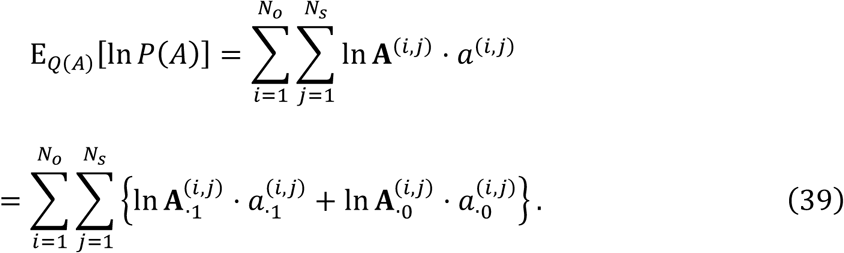

Furthermore, the synaptic update rules are derived from Equation (38) as

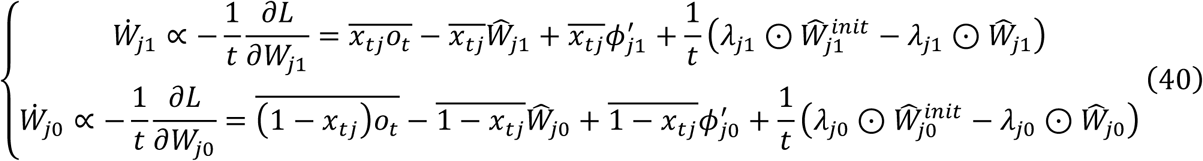

The fixed point of Equation (40) is provided as

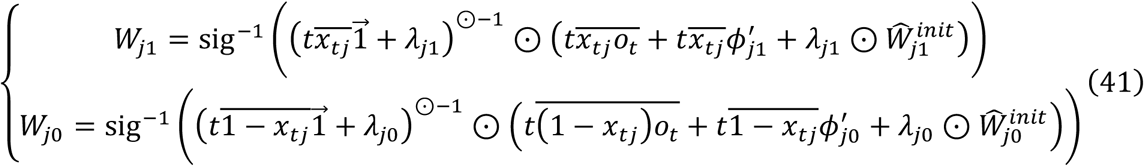

Note that the synaptic strengths at *t* = 0 are computed as 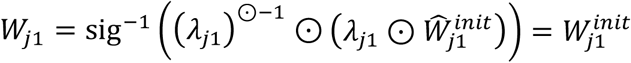. Again, one can see the formal correspondence between the final values of the synaptic strengths given by Equation (41) in the neural network formation and the parameter posterior given by Equation (8) in the variational Bayesian formation. As the Dirichlet parameter of the posterior 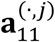 is decomposed into the outer product 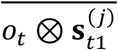 and the prior 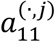, they are associated with 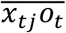 and 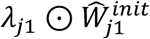, respectively. Thus, Equation (8) corresponds to Equation (41). Hence, for a given constant set 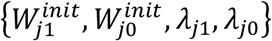, we identify the corresponding parameter prior 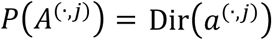, given by

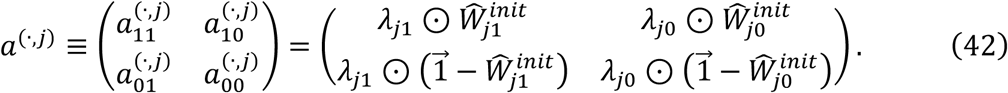

In summary, one can establish the formal correspondence between neural network and variational Bayesian formations, in terms of the cost functions (Equation (4) vs. Equation (38)), priors (Equation (18) and Equation (42)), and posteriors (Equation (8) vs. Equation (41)). This means that a neural network successively transforms priors *P*(*S*_*t*_), *P*(*A*) into posteriors *Q*(*S*_*t*_), *Q*(*A*), as parameterised with neural activity, and initial and final synaptic strengths (and thresholds). Crucially, when increasing number of observations, this process is asymptotically equivalent to that of variational Bayesian inference, under a specific likelihood function.

### Footnotes

1 Strictly speaking, the generative model used in this paper is a hidden Markov model (HMM) because we do not consider probabilistic transitions between hidden states that depend upon control variables. However, for consistency with the literature on variational treatments of discrete state space models, we retain the MDP formalism; noting that we are using a special case (with unstructured state transitions).

